# Annotated genome of the Eucalyptus snout beetle, *Gonipterus* sp. n. 2 (Coleoptera, Curculionidae)

**DOI:** 10.64898/2025.12.03.691750

**Authors:** Jade S. Ashmore, Gudrun Dittrich-Schröder, Rosa S. Knoppersen, Bernard Slippers, Almuth Hammerbacher, Tuan A. Duong

## Abstract

*Gonipterus* species, or Eucalyptus snout beetles, are defoliators that damage *Eucalyptus* trees in plantations globally. The *Gonipterus* sp. n. 2 genome was sequenced using both Oxford Nanopore and Illumina sequencing platforms which produced 76.41 Gb long-read and 57.1 Gb short-read sequence data, respectively. Genome assembly using these data resulted in 1,023 contigs, with an N50 of 2.78 Mb and a genome size of roughly 1.54 Gb. Genome completeness analysis using BUSCO resulted in a score of 98.2%. We used Braker3 to annotate the assembled *Gonipterus* sp. n. 2 genome using transcriptomic data from gut and reproductive tissues, as well as and available protein sequences from selected Coleoptera species. Genome annotation resulted in 42,343 protein coding gene models and a proteome BUSCO completeness score of 99%. Protein clustering with 13 other insect species using OrthoFinder identified 22,245 families with 1,398 families unique to *Gonipterus* sp. n. 2. The number of *cytochrome P450 monooxygenase* genes in *Gonipterus* sp. n. 2 (n = 119) was greater than the other 13 insect species used in comparison. The genome of *Gonipterus* sp. n. 2 will be a valuable resource to assist in unravelling various aspects of the weevil’s life history, such as the metabolism of xenobiotics or the production of pheromones, and to develop alternative pest control methods.

## 1.) Introduction

*Gonipterus* species, or Eucalyptus snout beetles, are coleopterans that damage *Eucalyptus* trees in plantations globally (Tooke 1955). International trade has resulted in the introduction of *Gonipterus* species from their native ranges into other *Eucalyptus*-growing countries, impacting *Eucalyptus* plantation forests globally (Wingfield *et al*. 2008; Hurley *et al*. 2016; SCHRÖDER *et al*. 2020). *Eucalyptus* defoliation by *Gonipterus scutellatus* ranges between 5% and 80% of plantation trees, depending on the season and the management techniques used (Loch and Matsuki 2010). Reliable and sustainable management strategies are desperately needed to manage this forestry pest.

With the establishment of next-generation sequencing, the amount of genomic data available for model and non-model organisms, including Coleoptera, has increased exponentially. To date, there are 331 and 840 coleopteran genomes available on InsectBase (https://www.insect-genome.com/genome) and National Centre for Biotechnology Information (NCBI) (https://www.ncbi.nlm.nih.gov/), respectively, with these coleopterans most commonly classified as pests (Wang *et al*. 2025). While InsectBase and NCBI contain 33 and 106 curculionid genomes, respectively, this study represents the first genome of a curculionid from the genus *Gonipterus* (Wang et al. 2025). Coleopteran genomes exhibit substantial size variation, ranging from just 18.22 Mb in the scarabid *Coptodactyla brooksi,* to 2714.43 Mb in the lycid *Platerodrilus igneus* (Wang *et al*. 2025).

The availability of coleopteran genomes has facilitated the exploration of a broad range of research questions. Specifically, Curculionidae genomes have been used to investigate topics such as host adaptation, characterisation of symbiotic relationships, identification of genes involved in environmental tolerance, characterisation of range expansion, microsatellite mining, identification of chemoreceptor genes, and to determine population structure (Apriyanto and Tambunan 2021; Navarro-Escalante *et al*. 2021; Keeling *et al*. 2022; MOHD Rodzik *et al*. 2023; Wang *et al*. 2023; Biswas *et al*. 2024; Chen *et al*. 2024). From an evolutionary perspective, research has also focused on protein evolution, such as that of luciferin, which is involved in bioluminescence in fireflies (Zhang *et al*. 2020). In addition, these coleopteran genomes also allow comparative genomic studies to inform the development of gene-based pest control methods (Chu *et al*. 2018).

Detoxification is a crucial adaptive mechanism for insect pests, as it enables them to neutralize harmful substances, such as plant secondary metabolites and insecticides, enhancing their survival, prevalence, and ability to spread. In insects, cytochrome P450 monooxygenases (P450s) play a key role in the metabolic detoxification of xenobiotics, such as plant secondary metabolites and insecticides (Feyereisen 2012; Cui *et al*. 2016; Lu *et al*. 2021). These enzymes reduce the biological activity of toxic compounds by adding an oxygen atom, thereby neutralizing their effects. The range of xenobiotics detoxified by insect P450s can vary from narrow to broad, while distantly related P450 enzymes may degrade the same compounds with differing efficiencies (Cui *et al*. 2016). Increased gene copy number or enhanced expression of P450 genes is often linked to greater resistance to insecticides (Feyereisen 2012). As a result, numerous studies have focused on identifying the specific P450 genes and gene superfamilies involved in detoxification, as well as their expression in response to insecticide exposure (Zhu *et al*. 2013; Evans *et al*. 2018; Zhang *et al*. 2023).

This article presents the first genome of *Gonipterus* sp. n. 2 (Coleoptera, Curculionidae), an important resource for better understanding the biology of this *Eucalyptus* pest and enabling further research into alternative pest control mechanisms.

## 2.) Results and Discussion

A total of 57.1 Gb of short-read DNA sequencing data (151 bp paired-end reads) and 23.4 Gb RNA sequencing data were generated with the Illumina HiSeq platform. Long read PromethION sequencing yielded 76.41 Gb data with read N50 of around 22 kb. Genome profiling using short reads with GenomeScope 2.0 resulted in an estimated genome size of 1.8 Gb. The primary NECAT assembly had 2,589 contigs, an N50 of 2.6 Mb and an assembled genome size of 1.76 Mb. After polishing and purging of haplotypes, the final haploid genome assembly had 1,023 contigs, an N50 of 2.78 Mb and a haplotype genome size of 1.54 Gb (Table 1). BUSCO analysis of the final assembly using the “insecta_odb10” dataset resulted in a completeness score of 98.3% (Manni *et al*. 2021).

**Table 1.** Statistics of the *Gonipterus* sp. n. 2 genome assembly and annotation.

| Genomic features |  |
| --- | --- |
| Genome assembly |  |
| Assembly size (bp) | 1 545 062 005 |
| Number of contigs | 1023 |
| N50 contig length (bp) | 2 782 955 |
| L50 | 163 |
| GC content | 32.24% |
| BUSCO (insecta_odb10, n=1367) | 98.3% |
| Gene models |  |
| Number of gene models | 42 343 |
| Mean gene length (bp) | 16 401 |
| Mean coding sequence length (bp) | 972 |
| Mean number of exons per gene | 3.4 |
| Mean exon length (bp) | 325 |
| Mean intron length (bp) | 6239 |
| BUSCO (insecta_odb10, n=1367) | 99% |

RepeatModeler identified 4,917 repeat families in the genome, comprising 73.46% of the whole genome when masked with RepeatMasker. The Braker3 pipeline predicted 42,343 protein-coding genes with BUSCO score of 99%, higher than the scores obtained from BUSCO run on the assembly, indicating that the annotation has sufficiently covered the organism’s gene space. Protein clustering of *Gonipterus* sp. n. 2 predicted proteome with 13 other insect species (11 Coleoptera, one Lepidoptera and one Neuroptera (Table 2)) using OrthoFinder resulted in 22,245 orthogroups. These species were selected based on the criterion that transcriptomic data were used as gene evidence during genome annotation, suggesting that their proteomes are of high quality. Of the identified orthogroups, 3,770 (16.95%) were shared by *Gonipterus* sp. n. 2, selected coleopteran species, and the two non-coleopteran species used as the outgroup (Figure 1). A total of 1,398 (6.28%) orthogroups were unique to *Gonipterus* sp. n. 2.

**Figure 1.**
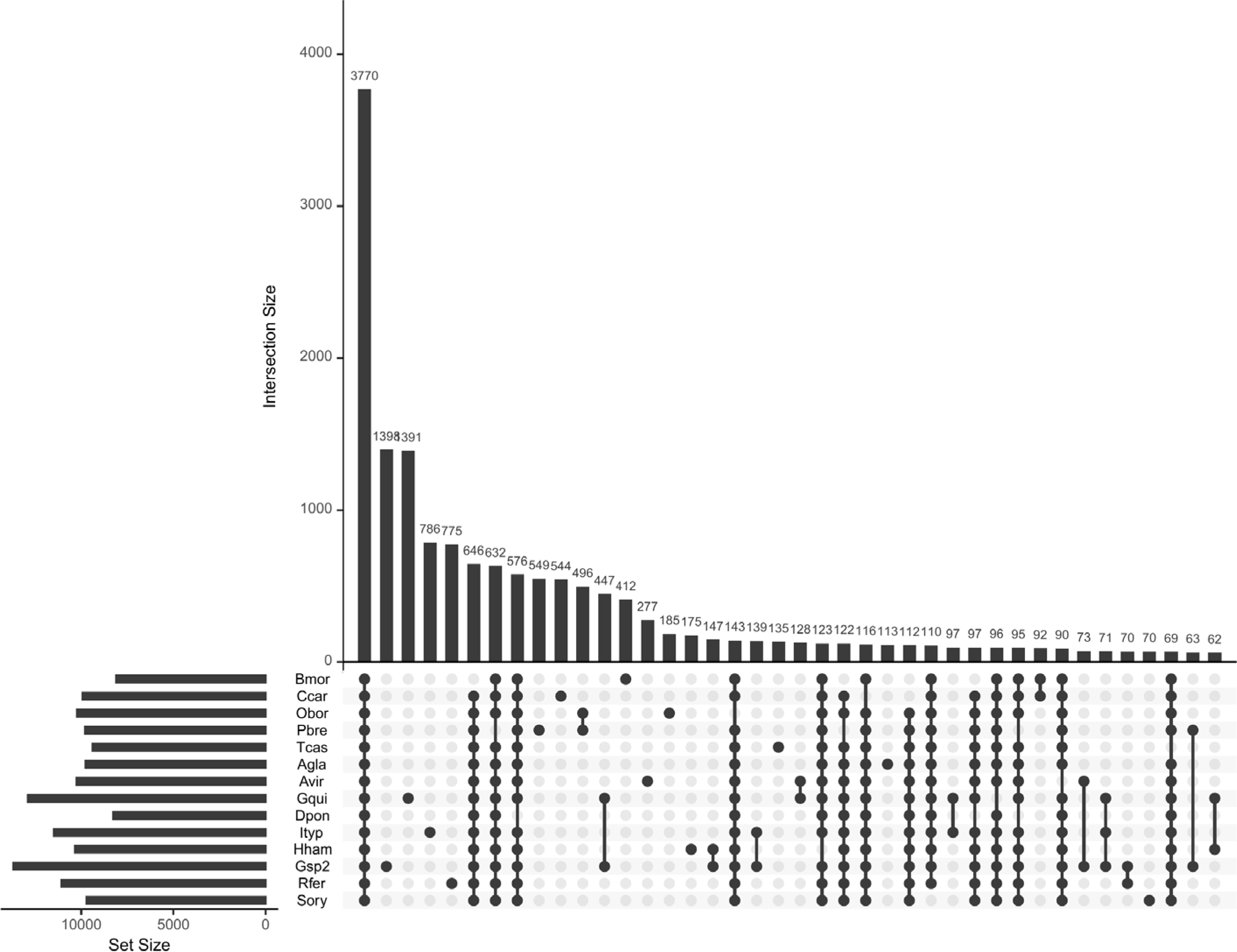
Upset-Plot of the number of gene families shared between the species and unique to the species. There were 3770 genes shared by all the species and the number of genes unique to an individual species was between 1398 and 70 genes by *Gonipterus* sp. n. 2 and *S. oryzae*, respectively. The set size for the species was between 5000 and 15000. The abbreviations for the species names are as follows Bmor was *B. mori*, Ccar was *C. carnea*, Obor was *Oryctes borbonicus* (Coleoptera, Scarabaeidae), Pbre was *Protaetia brevitarsis* (Coleoptera, Scarabaeidae), Tcas was *T. castaneum*, Agla was *Anoplophora glabripennis* (Coleoptera, Cerambycidae), Avir was *Altica viridicyanea* (Coleoptera, Chrysomelidae), Gqui was *Gonioctena quinquepunctata* (Coleoptera, Chrysomelidae), Dpon was *Dendroctonus ponderosae* (Coleoptera, Curculionidae), Ityp was *Ips typographus* (Coleoptera, Curculionidae), Hham was *Hypothenemus hampei* (Coleoptera, Curculionidae), Gsp2 was *Gonipterus* sp. n. 2 (Coleoptera, Curculionidae), Rfer was *Rhynchophorus ferrugineus* (Coleoptera, Curculionidae) and Sory was *S. oryzae*.

**Table 2.** A list of the selected coleopteran species and non-coleopteran species used during this study.

| **Insect species** | **Classification** |
| --- | --- |
| *Anoplophora glabripennis* | Coleoptera, Cerambycidae |
| *Altica viridicyanea* | Coleoptera, Chrysomelidae |
| *Bombyx mori*<sup>a</sup> | Lepidoptera, Bombycidae |
| *Chrysoperla carnea*<sup>a</sup> | Neuroptera, Chrysopidae |
| *Dendroctonus ponderosae* | Coleoptera, Curculionidae |
| *Gonioctena quinquepunctata* | Coleoptera, Chrysomelidae |
| *Hypothenemus hampei* | Coleoptera, Curculionidae |
| *Ips typographus* | Coleoptera, Curculionidae |
| *Oryctes borbonicus* | Coleoptera, Scarabaeidae |
| *Protaetia brevitarsis* | Coleoptera, Scarabaeidae |
| *Rhynchophorus ferrugineus* | Coleoptera, Curculionidae |
| *Sitophilus oryzae* | Coleoptera, Curculionidae |
| *Tribolium castaneum* | Coleoptera, Tenebrionidae |

Note: <sup>a</sup>, non-coleopteran insect species used as outgroups.

CAFÉ analysis of *Gonipterus* sp. n. 2 proteome together with those from 11 other coleopteran species, *Bombyx mori* (Lepidoptera, Bombycidae) and *Chrysoperla carnea* (Neuroptera, Chrysopidae) identified 437 gene families with significant expansion and 110 gene families with significant contraction (Figure 2). One hundred and nineteen P450 genes were found in 22 gene families in *Gonipterus* sp. n. 2, while the remaining 13 species had between 45 and 105 *cytochrome P450 monooxygenase* genes (Figure 2). Four *cytochrome P450 monooxygenase* gene families in *Gonipterus* sp. n. 2 experienced significant expansion and contained 79 (66.39%) of the identified *cytochrome P450 monooxygenase* genes (Figure 3.A to 3.D). None of the *cytochrome P450 monooxygenase* gene families in *Gonipterus* sp. n. 2 experienced significant contraction.

**Figure 2.**
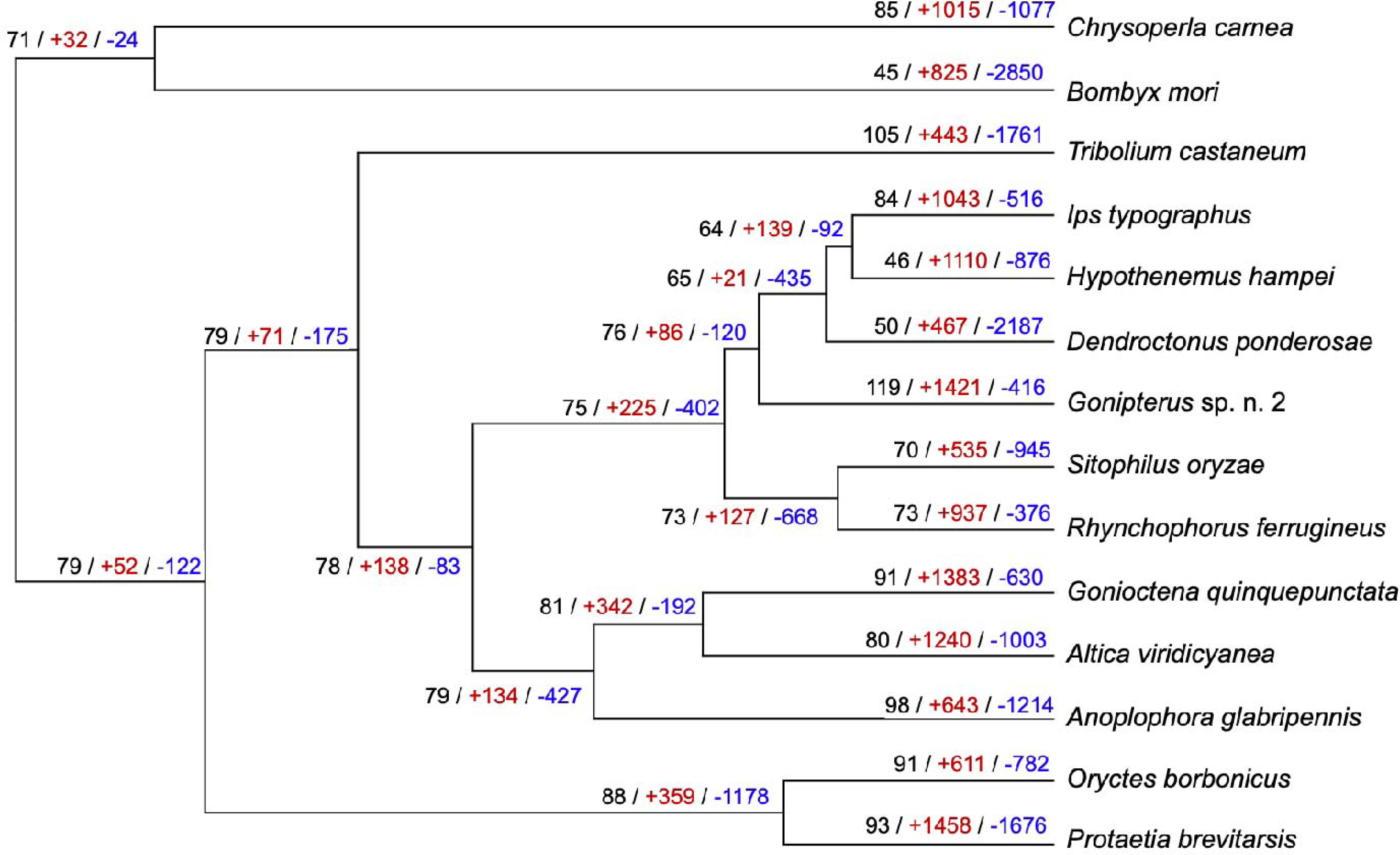
The number of *cytochrome P450 monooxygenase* genes identified and the number of gene families with significant expansion and contraction. The largest number of *cytochrome P450 monooxygenase* genes, indicated in black, was identified in *Gonipterus* sp. n. 2 (n = 119). The number of *cytochrome P450 monooxygenase* genes identified in selected coleopteran species ranged from 46 to 105 genes. *Chrysoperla carnea* and *B. mori* had 85 and 45 *cytochrome P450 monooxygenase* genes, respectively. The number of gene families with significant expansion and contraction is indicated in red and blue. *Gonipterus* sp. n. 2 has the largest significant expansion and smallest significant contraction of gene families of the selected coleopteran species and the outgroups.

**Figure 3:**
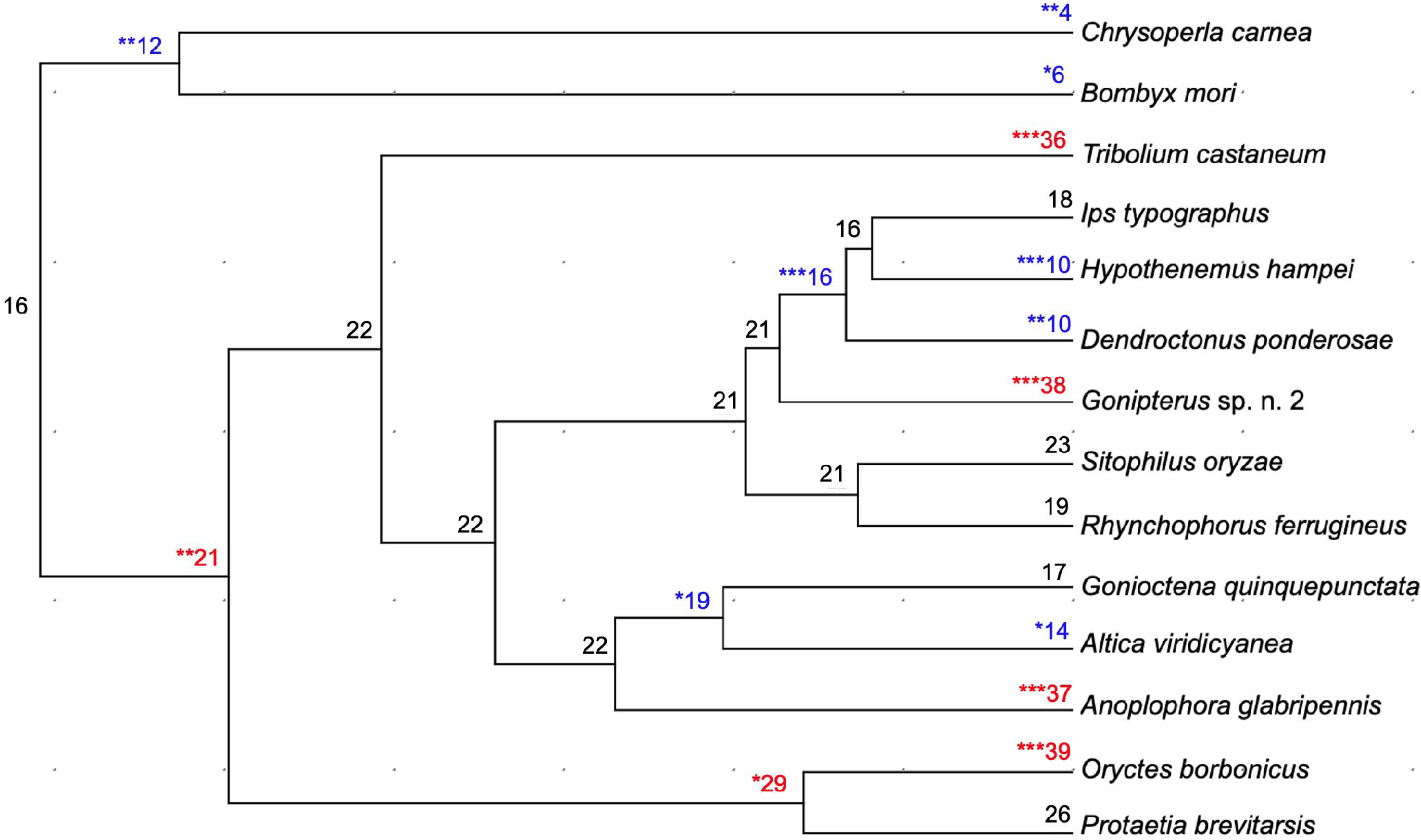
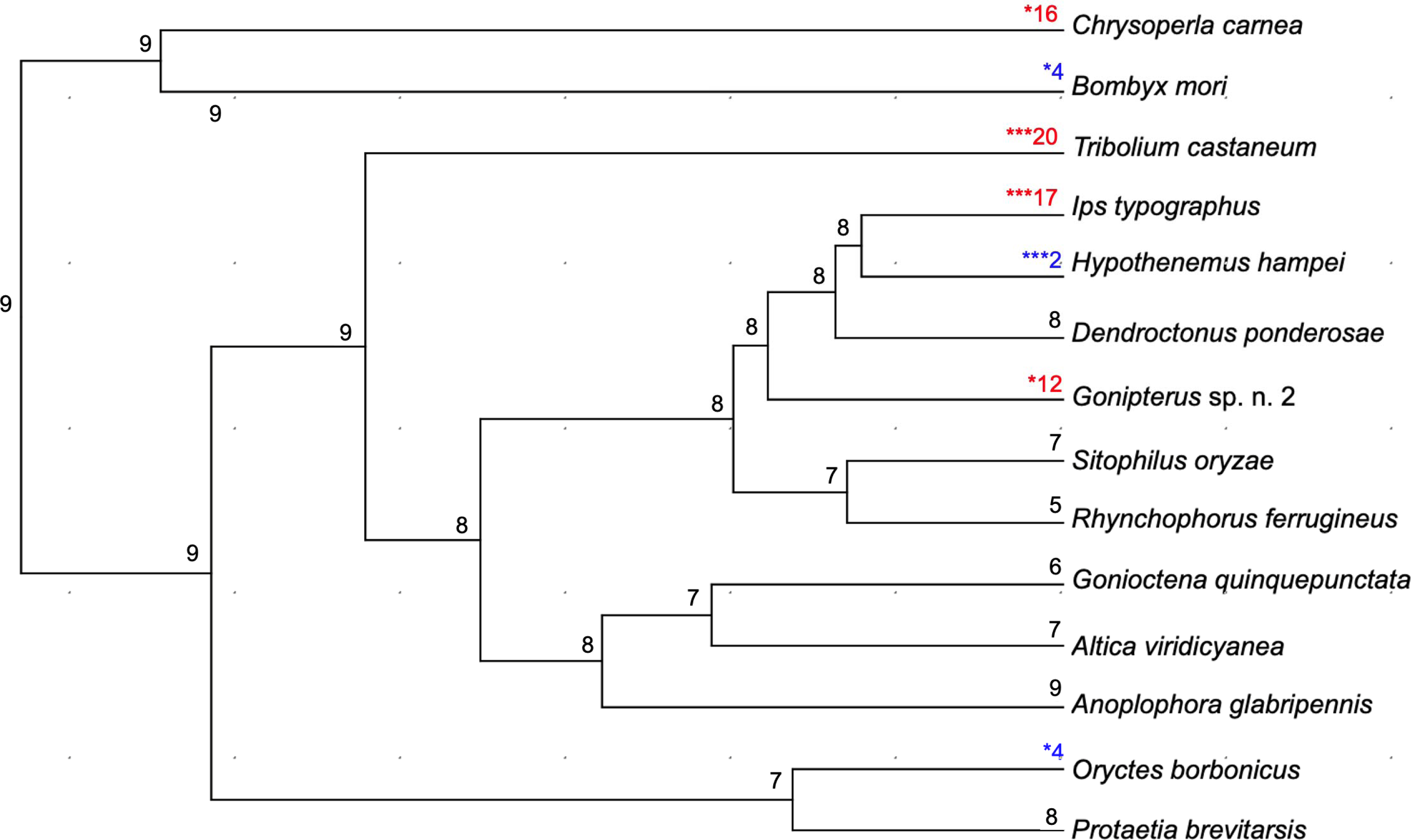
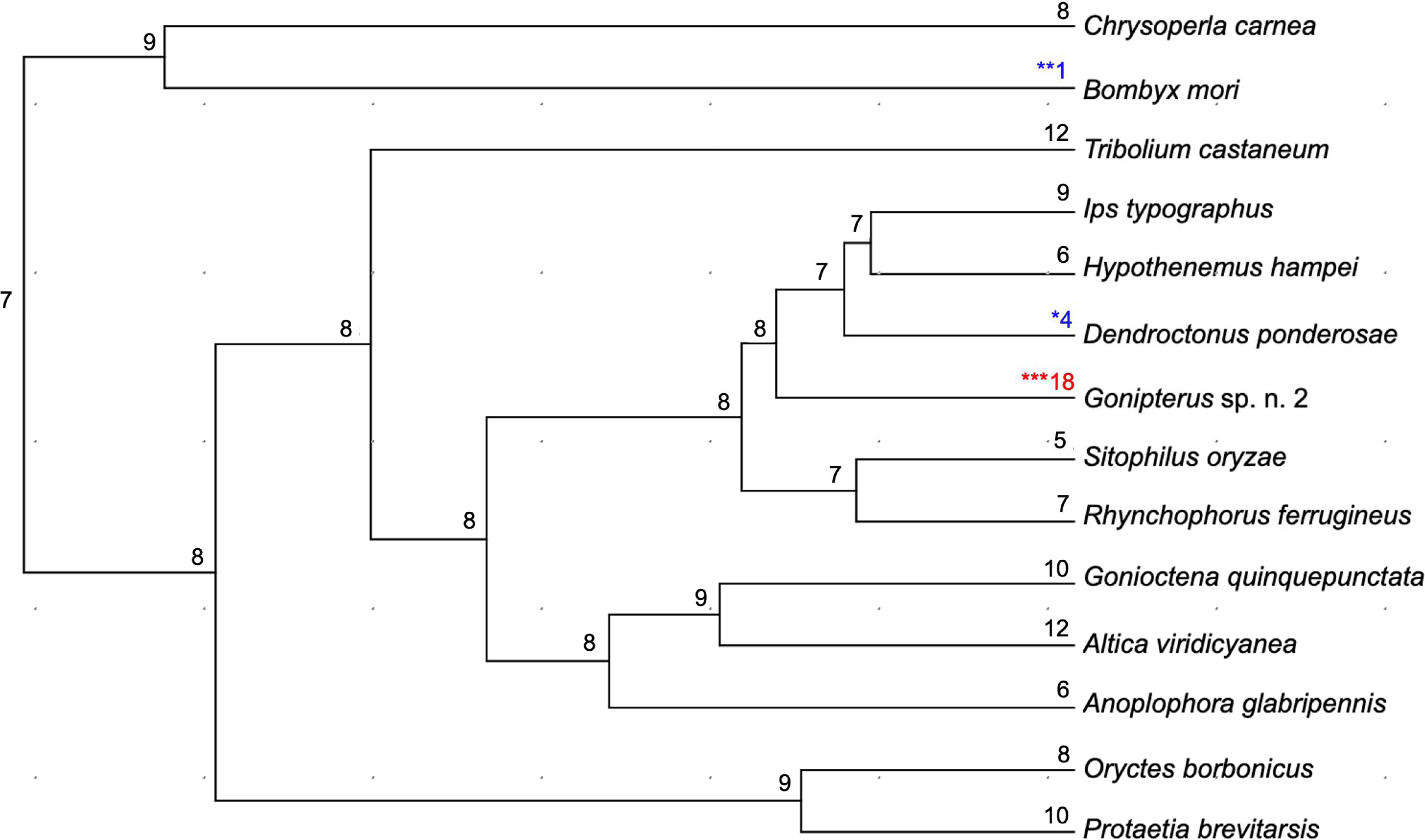
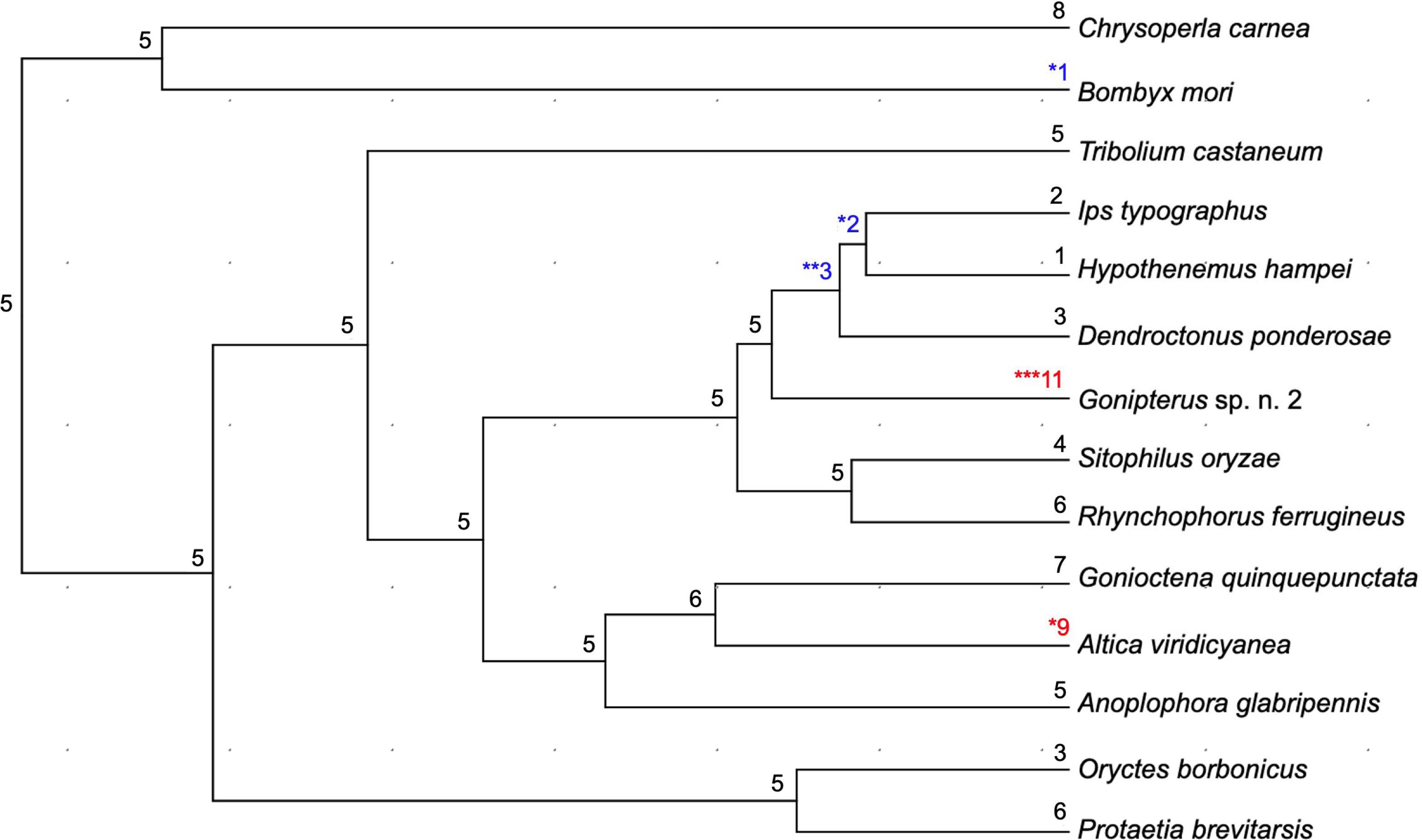
Phylogenetic trees illustrating the expansion and contraction of Orthogroups OG0000010 (A.), OG0000089 (B.), OG0000101 (C.) and OG0000244 (D.) in *Gonipterus* sp. n. 2, selected coleopteran species and the outgroups. The *cytochrome P450 monooxygenase* gene family protein domain was identified in Orthogroups by the Blast2GO PRO plug-in on the CLC Genomics Workbench. The numbers in red and blue at the nodes indicate expansion and contraction of the orthogroups. The numbers in black at the nodes indicate no expansion or contraction occurred. The statistical significance of the expansion or contraction at the individual nodes is indicated with asterisks; *, p < 0.05; **, p < 0.01; ***, p < 0.001.

*Gonipterus* sp. n. 2 had the highest number of *cytochrome P450 monooxygenase* genes (119 genes) amongst the species assessed. The expansion of *cytochrome P450 monooxygenase* genes may enable *Gonipterus* sp. n. 2 to detoxify the secondary metabolites in *Eucalyptus* leaves. *Eucalyptus* species have a high diversity of secondary metabolites with different biological activities, including insecticidal, antimicrobial, and antifeedant (BREZÁNI AND KAREL 2013; Danna *et al*. 2024). Insect-repellent and insecticidal activities of the *Eucalyptus* secondary metabolites were effective against different coleopterans, including *Tribolium castaneum* (Coleoptera, Tenebrionidae) and *Sitophilus oryzae* (Coleoptera, Curculionidae) (Danna *et al*. 2024). However, *Gonipterus* sp. n. 2 was attracted to the volatiles of damaged leaves from *Eucalyptus* host species (Bouwer *et al*. 2014). Increased diversity, copy number and expression of *cytochrome P450 monooxygenase* genes in insects is associated with increased xenobiotic resistance (Feyereisen 2012).

The *Gonipterus* sp. n. 2 genome may enable the development of alternative control methods, such as genetic pest control, to supplement currently applied pest control measures. Other coleopteran genomes have also been utilized to develop alternative pest control strategies. Specifically, these genomes have been leveraged to identify potential target genes for RNAi-based pest control and to facilitate CRISPR/Cas9 genome editing (Segers *et al*. 2023; Johny *et al*. 2024; Zhang *et al*. 2024). Such research in Coleoptera has primarily focused on model organisms, such as *Tribolium castaneum*, highly invasive species like *Harmonia axyridis* and insect pests of economic importance, such as *Leptinotarsa decemlineata* (Gui *et al*. 2020; Wu *et al*. 2022; Markley *et al*. 2024). Knowledge gained from the *Gonipterus* sp. n. 2 genome thus not only provides opportunities to direct future research to understand the beetle’s biology, but potentially also to improve and develop new pest control methods.

## 3.) Materials and Methods

### 3.1.) DNA extraction and genome sequencing

For long-read sequencing, *Gonipterus* sp. n. 2 adults were collected from *Eucalyptus* plantations near Greytown (KwaZulu-Natal, South Africa) (coordinates: 29.218415°S and 30.679624°E) during September 2021. The abdominal tissue only was used for DNA extraction following the Monarch High Molecular Weight (HMW) DNA Extraction Kit protocol for tissue, with slight modifications to the manufacturer’s instructions. The following modifications were made in Part 1: Tissue lysis. At step 5, the lysate mixture was incubated at 56 °C for 15 minutes with 1,150 rpm agitation, and, subsequently, 30 minutes without agitation to increase the DNA yield. During the 30 minutes incubation step without agitation, the lysate mixture was inverted every 5 minutes to ensure complete lysis. At step 9, the sample was centrifuged at 25 °C at 16,000 rcf for 15 minutes. At step 11, the DNA phase was aliquoted into a clean Eppendorf tube and centrifuged at 25 °C at 16,000 rcf for 15 minutes. After the second centrifugation step, the adipose was pipetted from the solution, and the DNA phase was transferred to a labelled Monarch 2 ml Tube. The DNA was separated from the solution and eluted following the HMW gDNA Binding and Elution procedure of the Monarch HMW DNA Extraction Kit protocol for tissue.

For short read sequencing, *Gonipterus* sp. n. 2 adults were collected from *Eucalyptus* plantations near Melmoth (KwaZulu-Natal, South Africa) (coordinates: 28.562856 °S, 31.191291 °E) during October 2020. The gut content was removed and DNA extraction was performed following the E.N.Z.A ® Insect DNA Kit (Omega Bio-tek). The samples were frozen in liquid nitrogen and ground to a fine powder. Proteinase K (25 μl) and CTL Buffer (350 μl) were mixed with the ground samples, and the mixture was incubated overnight at 37 °C. The DNA was isolated, and RNA was degraded following the E.N.Z.A ® Insect DNA Kit (Omega Bio-tek) protocol.

The *Gonipterus* sp. n. 2 genome was sequenced with the Illumina and Oxford Nanopore sequencing platforms. For Illumina sequencing, a paired-end library (350 bp median insert size) was prepared using TruSeq PCR-free protocol and sequenced on the HiSeq platform at Macrogen (Seoul, Korea) to obtain 151 bp paired-end reads. Read quality from the Illumina data was assessed with FastQC (v0.11.9) (https://www.bioinformatics.babraham.ac.uk/projects/fastqc/), and low-quality bases and remaining Illumina adaptors were removed with Trimmomatic (v0.38) (Bolger *et al*. 2014). For Nanopore sequencing, the library was constructed using the ligation sequencing kit (SQK-LSK110) and sequenced on the FLO-PRO002 flow cell (PromethION) at Centre for Genome Innovation at the University of Connecticut. Basecalling was conducted using Guppy (v5.1.12).

### 3.2.) RNA extraction and sequencing

For RNA sequencing, *Gonipterus* sp. n. 2 adults were collected from *Eucalyptus* species near Greytown (KwaZulu-Natal, South Africa) (coordinates: 29.1934795° S and 30.6082423° E) during January 2023. RNA extraction was performed using the gut tissue of the *Gonipterus* sp. n. 2 adults with the InviTrap® Spin Plant RNA Mini Kit (Invitek Diagnostics, Germany) following the manufacturer’s instructions. Briefly,10 samples, each containing the gut tissue from *Gonipterus* sp. n. 2 adults, were frozen in liquid nitrogen and ground into a fine powder. Cells were lysed by adding 900 μl of Lysis Solution RP. The mixture was incubated for 30 minutes at 55 °C and vortexed every five minutes. The DNA was removed by centrifuging the mixture at 16,160 rcf for one minute and filtering the resulting supernatant with a Prefilter. The RNA was bound to an RNA Spin Filter and washed following the manufacturer’s instructions. To elute the RNA, 30 μl of the Elution Buffer R was added to the RNA Spin Filter, incubated for two minutes at 25 °C, and then centrifuged for one minute at 11,000 rcf. The eluted RNA was immediately stored at −80 °C. A paired-end RNA library was constructed using the TruSeq Stranded mRNA Library Prep Kit, and the library was sequenced with Illumina HiSeq sequencing to obtain 151 paired-end reads at Macrogen Europe (The Netherlands). Adaptors and low-quality RNA sequences of reads were removed with Trimmomatic (v0.38).

### 3.3.) Genome assembly and annotation

The genome size and heterozygosity of *Gonipterus* sp. n. 2 were estimated from the trimmed Illumina data with JELLYFISH v.1.1.12 (MARÇAIS AND KINGSFORD 2011) and GenomeScope 2.0 (Ranallo-Benavidez *et al*. 2020), with a k-mer value of 21. The NECAT pipeline (Chen *et al*. 2021) was used to assemble the uncorrected reads from the Nanopore data using the estimated genome size obtained from GenomeScope and a minimum read length of 3 kb. The raw Nanopore data were mapped to NECAT assembly with minimap 2.0 (Li 2018), and the mapping file was used to polish the assembled genome with racon v1.3.1 (Vaser *et al*. 2017) for three iterations. The trimmed Illumina data were mapped to the racon-polished genome with BWA v0.7.17 (Vasimuddin *et al*. 2019) and used to further polished the genome with Pilon v1.23 (Walker *et al*. 2014) for three iterations. A final round of polishing was conducted with racon using trimmed Illumina data mapped to the Pilon-polished assembly. Haplotypes in the primary polished assembly were removed with Purge Haplotigs (Roach *et al*. 2018). The completeness of the polished genome was assessed with the Benchmarking Universal Single-Copy Orthologs (BUSCO) v.4.0.5 utilising the “insecta_odb10” dataset (Manni *et al*. 2021).

Structural annotation was carried out using the Braker3 pipeline (Stanke *et al*. 2006; Stanke *et al*. 2008; Buchfink *et al*. 2015; Hoff *et al*. 2019; BRŮNA *et al*. 2021), using Augustus as the gene predictor. RepeatModeler v2.0.2 (http://www.repeatmasker.org/RepeatModeler/) was used to construct a *de novo* repeat library, which was used to soft-mask the genome with RepeatMasker v 4.1.2 (http://www.repeatmasker.org/RepeatModeler/). RNA sequencing data from *Gonipterus* sp. n. 2 reproductive tissue (NCBI accession number: SAMN19700001), alimentary tissue (NCBI accession number: SAMN19700000) (Souza *et al*. 2022), and the trimmed RNA sequencing data from the gut of *Gonipterus* sp. n. 2 (PRJNA1189815) were aligned to the masked *Gonipterus* sp. n. 2 draft genome with HISAT2 v2.2.1 (Kim *et al*. 2019). Proteomes of the selected coleopteran species (Table 2) were mapped to the *Gonipterus* sp. n. 2 genome with GenomeThreader v1.7.4 (Gremme *et al*. 2005). The aligned transcriptomes and proteomes were used as evidences to train the *ab initio* prediction tool Augustus for the Braker3 pipeline. The Blast2GO PRO plug-in v1.20.14 on the CLC Genomics Workbench v20.0.4 was used to assign functions to the predicted proteins and identify P450 genes using the protein domain (Pfam = 00067).

### 3.4.) Analysis of the expansion of the cytochrome P450 gene family

Orthologous gene families from the protein sequences of *Gonipterus* sp. n. 2, selected coleopteran species (Table 2), *Bombyx mori* (Lepidoptera, Bombycidae) and *Chrysoperla carnea* (Neuroptera, Chrysopidae) were determined using OrthoFinder v2.5.5 (Emms and Kelly 2015), with the Diamond in sensitive mode (-S) and the inflation factor of 1.5. The coleopteran species and outgroups were selected from InsectBase (Accessed in June 2023) (Mei *et al*. 2022) based on the availability of genomes annotated with transcriptomic data, which enabled the prediction of a higher quality and more comprehensive protein set (Chen *et al*. 2017). To construct a species phylogeny, protein sequences of the single-copy orthologs were aligned with muscle v3.8.31 (Edgar 2022), and trees were inferred with IQ-TREE v 2.1.2 (Kalyaanamoorthy *et al*. 2017). A species phylogeny was inferred from individual gene trees with ASTRAL v5.7.7 (Zhang *et al*. 2018). The obtained phylogeny was rooted in *B. mori* and *C. carnea*, and the branch length was optimised with RAxML v8.2.11 (Stamatakis 2014) using the concatenate alignment of all single-copy genes. The rooted phylogenetic tree was transformed into an ultrametric tree and used as input in CAFÉ. CAFÉ v5 (Mendes *et al*. 2020) was used to assess the expansion and contraction of P450 gene families.

## Data availability

The assembled genome *Gonipterus* sp. n. 2 has been deposited in NCBI (https://www.ncbi.nlm.nih.gov/) under the accession number PRJNA1203571. The raw RNA sequencing data was deposited in NCBI with the accession number PRJNA1189815.

## CRediT authorship contribution statement

**Jade S. Ashmore**: Data Curation; Formal Analysis; Investigation; Resources; Writing – Original Draft Preparation; Writing – Review & Editing. **Gudrun Dittrich-Schröder**: Conceptualization; Funding Acquisition; Investigation; Project Administration; Resources; Supervision; Writing – Original Draft Preparation; Writing – Review & Editing. **Rosa S. Knoppersen**: Data Curation; Resources; Writing – Review & Editing. **Bernard Slippers**: Conceptualization; Project Administration; Supervision; Writing – Review & Editing. **Almuth Hammerbacher**: Funding Acquisition; Resources; Writing – Review & Editing. **Tuan A. Duong**: Conceptualization; Data curation; Formal Analysis; Investigation; Project Administration; Supervision; Validation; Writing – Original Draft Preparation; Writing – Review & Editing.

## Acknowledgements

We would like to thank Mr Preston L. Shaw (FABI, University of Pretoria) for assisting with the genome submission process.

